# Differential impact of transdiagnostic, dimensional psychopathology on multiple scales of functional connectivity

**DOI:** 10.1101/2021.03.05.434151

**Authors:** Darsol Seok, Joanne Beer, Marc Jaskir, Nathan Smyk, Adna Jaganjac, Walid Makhoul, Philip Cook, Mark Elliott, Russell Shinohara, Yvette I. Sheline

**Affiliations:** Center for Neuromodulation in Depression and Stress, Department of Psychiatry, Perelman School of Medicine, University of Pennsylvania, United States; Penn Statistics in Imaging and Visualization Center, Department of Biostatistics, Epidemiology, and Informatics, Perelman School of Medicine, University of Pennsylvania, United States; Department of Radiology, Perelman School of Medicine, University of Pennsylvania, United States; Department of Psychiatry, Perelman School of Medicine, University of Pennsylvania, United States

## Abstract

**Introduction:** Dimensional psychopathology strives to associate different domains of cognitive dysfunction with brain circuitry. Connectivity patterns as measured by functional magnetic resonance imaging (fMRI) exist at multiple scales, with global networks of connectivity composed of microscale interactions between individual nodes. It remains unclear how separate dimensions of psychopathology might differentially impact these different scales of organization.

**Methods:** Patients experiencing anxious misery symptomology (depression, anxiety and trauma; n = 192) were assessed for symptomology and received resting-state fMRI scans. Three modeling approaches (seed-based correlation analysis [SCA], support vector regression [SVR] and Brain Basis Set Modeling [BSS]), each relying on increasingly dense representations of functional connectivity patterns, were used to associate connectivity patterns with six different dimensions of psychopathology: anxiety sensitivity, anxious arousal, rumination, anhedonia, insomnia and negative affect. Importantly, a full 50 patients were held-out in a testing dataset, leaving 142 patients as training data.

**Results:** Different symptom dimensions were best modeled by different scales of brain connectivity: anhedonia and anxiety sensitivity were best modeled with single connections (SCA), insomnia and anxious arousal by mesoscale patterns (SVR) and negative affect and ruminative thought by broad, cortex-spanning patterns (BBS). Dysfunction within the default mode network was implicated in all symptom dimensions that were best modeled by multivariate models.

**Conclusion:** These results suggest that symptom dimensions differ in the degree to which they impact different scales of brain organization. In addition to advancing our basic understanding of transdiagnostic psychopathology, this has implications for the translation of basic research paradigms to human disorders.

## Introduction

Traditionally, mental illnesses have been conceptualized as disorder classes diagnosed based on the types, numbers and severity of symptoms reported by patients, as embodied in the Diagnostic and Statistical Manual of Mental Disorders (DSM). The DSM’s observational taxonomy persists in part due to a lack of reliable biomarkers for psychiatric disorders and remains the standard of practice for clinicians today. However, notable issues with the DSM system such as heterogeneity within disorder classes, high comorbidity between disorders and a lack of understanding of underlying mechanisms have motivated the development of an alternative framework: the National Institute of Mental Health’s (NIMH) Research Domain Criteria framework (RDoC;[1]). Whereas the DSM grouped diverse symptoms together based on common co-occurrence, RDoC delineates these symptoms into distinct dimensions of behavioral and cognitive (dys)function, with the goal of mapping these dimensions onto genetic, molecular, cellular and circuit mechanisms. The current study applies this dimensional approach to a class of disorders termed disorders of “anxious misery” [2, 3], which includes DSM diagnoses like generalized anxiety disorder (GAD), major depressive disorder (MDD), dysthymic disorder, and post-traumatic stress disorder (PTSD). Combined, these disorders affect over 800 million people worldwide and constitute the leading cause of disability [4]. Patients with these disorders exhibit dysfunction along various dimensions of neurocognitive functioning (constructs in the RDoC) such as fear, anxiety, frustrative nonreward, sleep-wakefulness and loss.

A key element for investigating these dimensions is the identification of their underlying neural circuits, part of NIMH’s goal to “map the connectomes for mental illness” (NIMH Strategic Plan, Strategy 1.3, Objective 1). Resting-state networks [5, 6] can be probed using noninvasive brain imaging to visualize the functional coactivation of specific regions of the brain, and previous work has implicated various patterns of hypo- and hyperconnectivity to anxious misery etiology [7]. Importantly, functional brain connectivity patterns exist at multiple scales [8-10], with large-scale patterns of connectivity composed of microscale interactions between individual functional nodes of the brain and mesoscale interactions between coherent communities of nodes. Different dimensions of symptomology may be best represented at different scales of brain connectivity; dysfunction in a single circuit may underlie the etiology of some dimensions of psychopathology while broader, global patterns of connectivity may best model other dimensions. Previous work on multiscale brain connectomics in psychiatry has focused on single-diagnosis, DSM-derived cohorts of patients [11, 12], and it remains unclear how transdiagnostic dimensions of psychopathology might differentially impact different scales of connectomics [12].

Different modeling approaches utilize information at different scales of functional connectomics. At one extreme are univariate seed-based correlation (SCA) approaches, which examine associations between behavioral variables and connectivity between just two regions, the selection of which is often determined *a priori*. In contrast, data-driven models such as support vector regression (SVR; [13]) and connectome-based predictive modeling (CPM; [14]) typically select a small subset of functional connections across the brain to generate multivariate symptom-brain associations. Finally, recent methods like brain basis set modeling (BBS; [15]) capitalize on connectivity patterns that span the entire brain, utilizing dimensionality reduction approaches like principal component analysis (PCA) to efficiently summarize large-scale patterns. It remains an open question which modeling approaches are best suited for particular dimensions of anxious misery symptomology.

An important consideration in assessing the generalizability of symptom-brain associations is the ability to replicate performance in a held-out testing dataset [16]. Systematic examinations of univariate and multivariate brain-behavior associations have revealed that samples far larger than those typically used in neuroimaging studies of psychopathology may be required to generate stable associations [17]. Further, notable examples of failed replications [18, 19] have revealed the perils of fitting and interpreting multivariate models, particularly when models rely on a small subset of functional connections selected through data-driven methods. Many of these attempts at replication stemmed from findings that incorrectly applied cross-validation, which requires complete encapsulation of information between folds in order to generate a reasonable estimate of out-of-sample performance [20]. Direct validation of model performance in a held-out sample is therefore an important step to assessing the potential generalizability of any symptom-brain association.

To this end, the present work examined transdiagnostic associations between six different dimensions of anxious misery symptomology (anxiety sensitivity, anxious arousal, ruminative thought, anhedonia, insomnia and negative affect) and functional connectivity, using three different modeling approaches. Importantly, the data were randomly divided into a training set and a held-out testing set to validate model performance.

## Materials and methods

### Sample characteristics

For a complete description of sample characteristics, screening procedures and participant assessment, readers are directed to the reference publication for this dataset [21].

194 participants experiencing symptoms of anxious misery (AM) and 48 healthy comparators (HC) were recruited through the community. Demographic information is summarized in Table 1, with additional demographic information in Supplemental Table 1. Two AM participants were removed from analyses due to excessive in-scanner motion, so we present demographics for 192 AM and 48 HC participants.

**Table 1.** Participant characteristics. Additional demographic information can be found in Supplemental Table 1.

| % Total | Category | Total<br>(N = 240) | Control<br>(N = 48) | Anxious Misery<br>(N = 192) |
| --- | --- | --- | --- | --- |
| **Sex** |  |  |  |  |
| 70.0% | Female | 168 | 34 | 134 |
| 30.0% | Male | 72 | 14 | 58 |
| **Age (median = 26)** |  |  |  |  |
| 28.3% | 18-23 y/o | 68 | 11 | 57 |
| 37.9% | 24-29 y/o | 91 | 23 | 68 |
| 17.5% | 30-35 y/o | 42 | 9 | 33 |
| 12.5% | 36-41 y/o | 30 | 4 | 26 |
| 1.3% | 42-47 y/o | 3 | 1 | 2 |
| 0.8% | 48-53 y/o | 2 | 0 | 2 |
| 1.7% | 54-59 y/o | 4 | 0 | 4 |
| **Race** |  |  |  |  |
| 10.4% | Asian | 25 | 6 | 19 |
| 18.8% | Black | 45 | 9 | 36 |
| 5.4% | Multiracial | 13 | 2 | 11 |
| 2.1% | Other | 5 | 2 | 3 |
| 3.3% | Undisclosed | 8 | 0 | 8 |
| 60.0% | White | 144 | 29 | 115 |
| **Ethnicity** |  |  |  |  |
| 7.1% | Hispanic | 17 | 3 | 14 |
| 91.2% | Not Hispanic | 219 | 45 | 174 |
| 1.7% | Undisclosed | 4 | 0 | 4 |
| **Medication status** |  |  |  |  |
| 25.6% | Medicated | - | - | 51 |
| 73.4% | Unmedicated | - | - | 141 |
| **Primary DSM-5 Diagnosis** |  |  |  |  |
| 47.4% | Major Depressive Disorder | - | - | 91 |
| 21.9% | Generalized Anxiety Disorder | - | - | 42 |
| 20.3% | Posttraumatic Stress Disorder | - | - | 39 |
| 6.77% | Persistent Depressive Disorder | - | - | 13 |
| 3.12% | Social Anxiety Disorder | - | - | 6 |
| 0.52% | Cyclothymic Disorder | - | - | 1 |

To ensure a truly transdiagnostic dataset, instead of using diagnosis from the Structure Clinical Interview for DSM-5 (SCID-5) as an inclusion criterion, participants were deemed eligible if their Neuroticism score on the Neuroticism-Extraversion-Openness Five-Factor Inventory (NEO FFI) was greater than one standard deviation above the population mean [22].

We did not require participants to cease taking psychotropic medication that they were currently taking. The majority of our study participants (73.4%) were unmedicated.

### Participant assessment

In order to assess participants’ symptom profiles, seven clinician-administered and self-report scales measuring a broad range of depressive and anxious symptomology were administered to all participants (see full list of assessments in Supplemental Methods and [21]).

### Hierarchical clustering of symptoms

To derive data-driven symptom communities from our assessment data, we applied a hierarchical clustering algorithm [23] on the Pearson correlation matrix of the 113 constituent items of our seven instruments, using only data from the AM group. After selecting an appropriate resolution (see Supplemental Methods for details), we derived symptom dimension scores for each participant by computing the mean across items belonging to each symptom community, after z-scoring.

### Imaging acquisition

Participants received T1w and four resting state fMRI scans (total acquisition time = 23:01) after their symptom assessments. Full imaging parameters and image acquisition protocols are detailed elsewhere [21].

### Image preprocessing

For full preprocessing details, readers are directed to the reference publication for this dataset [21]. 2 participants, both of whom were AM participants (0.83% of all participants) were removed from resting state analyses due to excessive in-scanner movement.

### Overview of modeling approaches

We tested the ability of three different modeling approaches, each relying on increasingly dense representations of functional connectomics, to generate robust, generalizable associations between imaging data and symptom dimensions. An overview of each of these modeling approaches is provided with complete details provided in the Supplemental Methods.

### Modeling Approach 1

#### Seed-based correlation analysis

Seed-based correlation analysis (SCA) associates behavioral variables (in our case, symptom dimension scores) with connectivity between a seed region and another region. Each symptom dimension was assigned specific seeds and masks based on extant literature (Table 2). Exploratory whole-brain analyses were also conducted. In cases where neither the initially proposed set of masks nor the whole-brain analyses generated a significant cluster, post-hoc analyses were conducted where further masks were examined (this was only necessary for the negative affect cluster, and only two additional sets of masks were examined post-hoc, see Table 2).

**Table 2.**
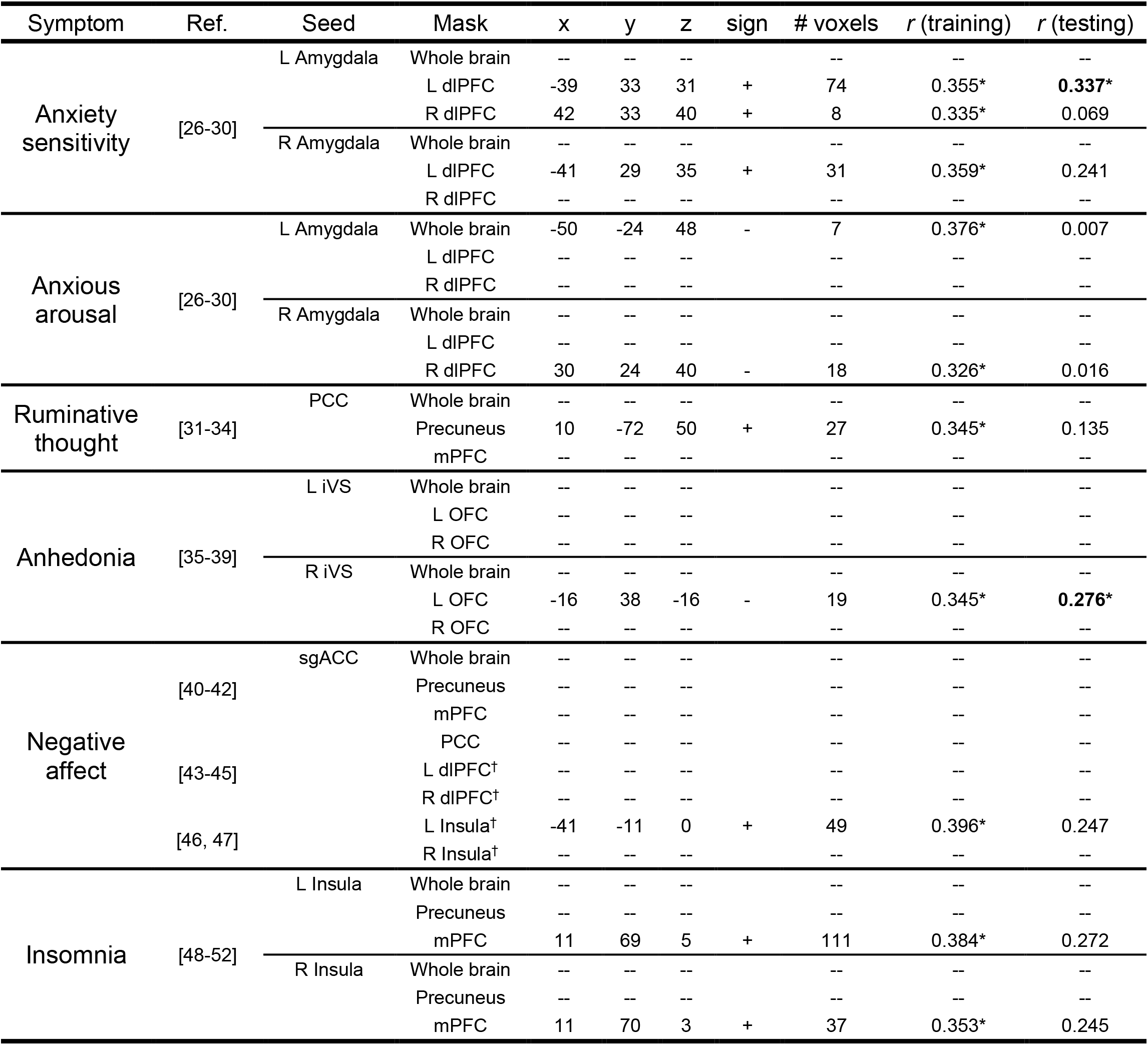
Seed-based correlation analysis results. Unique seeds and masks for each symptom cluster (in addition to a whole brain analysis) were assigned based on findings from extant literature (Ref.). Significant clusters were identified using a nonparametric permutation testing technique incorporating threshold-free cluster enhancement (implemented in FSL’s randomise; FWER < 0.05). Coordinates for cluster center of mass are in MNI space. “Sign” indicates the direction of association: “+” indicates that hyperconnectivity was associated with worse symptoms, while “-” indicates that hypoconnectivity was associated with worse symptoms. *r* (training) and *r* (testing) indicate Pearson correlations between observed and predicted symptom scores using a linear model using connectivity *z*-scores. * indicates correlation was significant at p < 0.05 using permutation testing. † indicates masks that were added post-hoc to identify a significant association in the training set. dlPFC = dorsolateral prefrontal cortex, PCC = posterior cingulate cortex, mPFC = medial prefrontal cortex, iVS = inferior ventral striatum, OFC = orbitofrontal cortex, sgACC = subgenual anterior cingulate cortex.

Clusters that exhibited significant associations based on permutation testing (*p* < 0.05) are reported. Linear models predicting symptom severity using connectivity were constructed using mean connectivity between the seed region and the identified cluster for each subject.

## Multivariate methods

The next two modeling approaches used the Power264 atlas, a well-validated parcellation that identifies 264 functional nodes of the brain [24]. In addition to these 264 nodes, we supplemented 14 additional nodes relevant to anxious misery disorders, adopted from the additional nodes used in [25] (see full table of additional seeds in Supplemental Table 3).

**Table 3.** Pearson’s correlations between actual and predicted symptom constructs in the held-out testing set using three different modeling approaches. In parentheses are permutation-based p-values where rows of imaging data were shuffled relative to symptom data in the training set and used to predict testing set symptom data (repeated 5000 times for each symptom and modeling approach). Also listed are the number of functional connectivity features (# of conn.) used for each model. Highlighted in **bold** are the best performing model approaches for each symptom construct. For SCA, the best performing association for each symptom was selected. * indicates correlation was significant at p < 0.05. SCA = seed-based correlation analysis. SVR = support vector regression. BBS = Brain Basis Set Modeling.

| Symptom | SCA |  | SVR |  | BBS |  |
| --- | --- | --- | --- | --- | --- | --- |
| | $r_{\text{testing}}$ ($p$) | # of conn. | $r_{\text{testing}}$ ($p$) | # of conn. | $r_{\text{testing}}$ ($p$) | # of conn. |
| Anxiety sensitivity | **0.337\* (0.017)** | **1** | 0.225 (0.062) | 500 | 0.284* (0.026) | 38503 |
| Anhedonia | **0.276\* (0.049)** | **1** | 0.185 (0.097) | 500 | 0.198 (0.086) | 38503 |
| Insomnia | 0.272 (0.058) | 1 | **0.344\* (0.006)** | **4600** | 0.313* (0.013) | 38503 |
| Anxious arousal | 0.016 (0.909) | 1 | **0.336\* (0.009)** | **1200** | 0.255* (0.046) | 38503 |
| Negative affect | 0.247 (0.088) | 1 | 0.016 (0.449) | 140 | **0.323\* (0.013)** | **38503** |
| Ruminative thought | 0.135 (0.351) | 1 | 0.167 (0.136) | 3000 | **0.313\* (0.014)** | **38503** |

For each of the following methods, we extracted the mean time series of each node using spherical ROIs (5mm radius) and then computed the Pearson correlation between each time series to generate a 278 × 278 functional connectivity matrix for each scan (38503 unique connections). Because participants completed four scans, the four resulting matrices were averaged to generate one matrix per participant.

### Modeling Approach 2

#### Support vector regression (SVR)

Support vector machines (SVMs) are a class of supervised machine learning techniques that attempt to learn a hyperplane that maximizes the margin between two classes, and extensions of this technique allow for continuous regression (SVR). Our cross-validated feature selection procedure indicated that SVR models performed best with a small subset of connectivity features (∼5% of the full matrix).

### Modeling Approach 3

#### Brain Basis Set Modeling (BBS)

Brain Basis Set Modeling is a recently developed technique that associates symptoms with a low dimensionality representation of global functional connectivity patterns [15]. In short, principal components analysis was used to generate a low-dimensional representation of global connectivity patterns, and then linear regression was used to associate each symptom dimension with connectivity component scores. Of the three modeling approaches, only BBS utilized data from HC participants (Supplemental Methods).

### Modeling assessment framework

Each of the models that was tested underwent the following assessment framework (Fig. 1) designed to test the performance of our association models on new, unseen data. First, age, sex and motion (quantified using mean RMS) were regressed from connectivity matrices, and age and sex were regressed from symptom scales. Second, while matching for age, sex and symptom severity, a random subset of 50 AM participants (26.0% of AM participants) were segregated to a testing set, leaving 142 AM participants for our training set. Next, within the training set, five-fold cross validation was employed to perform hyperparameter tuning and feature selection (Supplemental Methods) for the multivariate methods. Finally, model performance was assessed in the held-out testing set, again by computing the Pearson correlation between the predicted and actual symptom scores. Significance was determined using permutation testing (Supplemental Methods).

**Figure 1.**
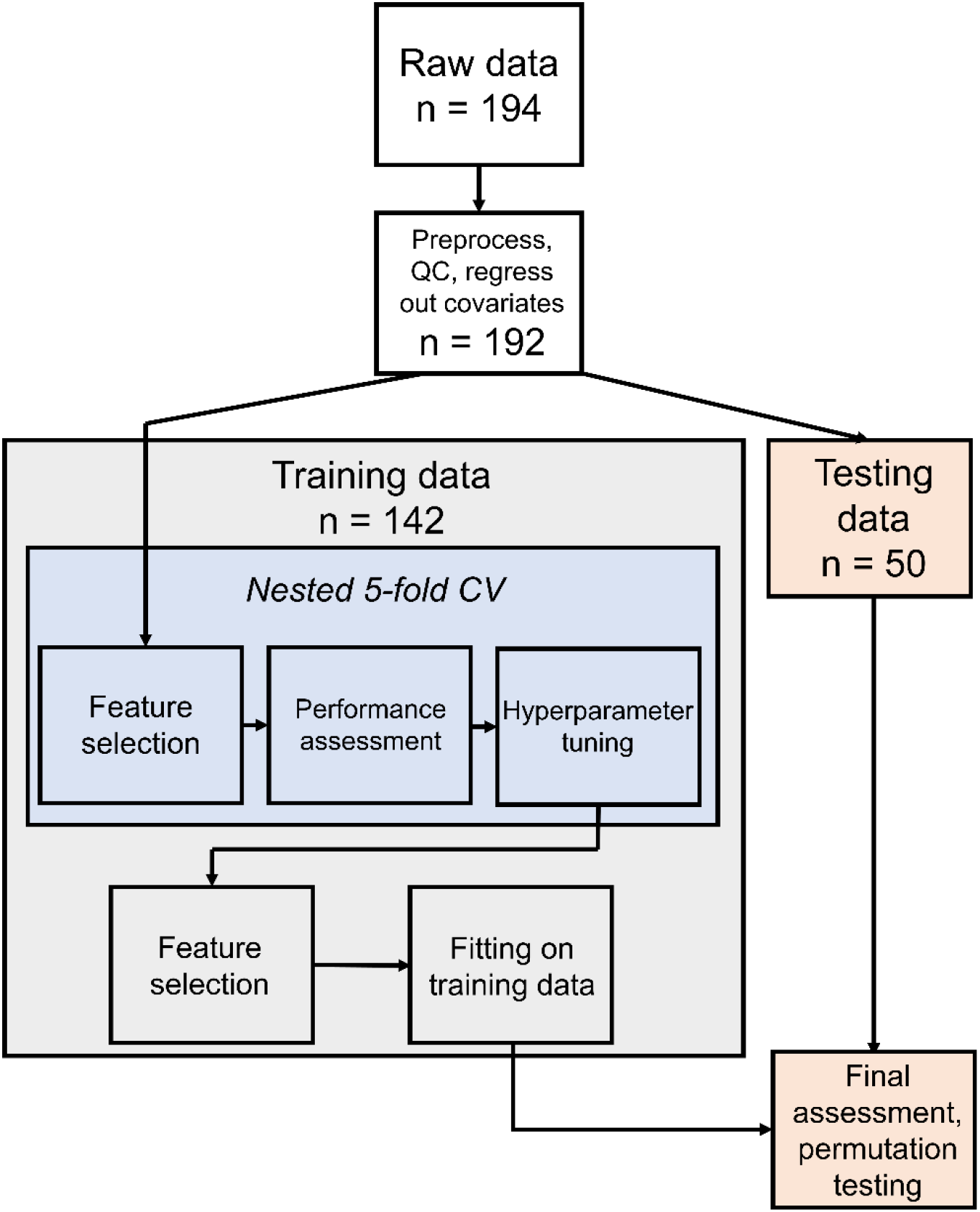
Flowchart of model assessment framework. After data are split into training and testing partitions, hyperparameter tuning in training data is accomplished using nested 5-fold cross validation. The best performing hyperparameters are used in a final model that is fit on the full training dataset. Finally, model performance is assessed using the testing data, and significance is determined using permutation testing. This process was repeated for each symptom dimension, each time using the same training/testing partition.

## Results

### Symptom clustering

Inspection of the symptom hierarchy revealed that items clustered according to clinically relevant dimensions of dysfunction (see Supplemental Figure 1 for complete symptom hierarchy). Symptoms first separated into two large communities, one with predominantly depressive symptoms and the other with largely anxiety-related symptoms. At finer resolutions, these two larger communities separated into three subcommunities each, resulting in six total communities (Fig. 2). Anxiety-related symptoms divided into the following communities: anxiety sensitivity (ASI item #3: “It scares me when my heart beats rapidly.”), anxious arousal (MASQ item #16: “Hands were shaky”) and ruminative thought (RTS item #4: “I can’t stop thinking about some things.”). Depression-related symptoms divided into the following communities: anhedonia (SHAPS item #3: “I would find pleasure in my hobbies and pastimes.”), insomnia (ISI item #1: “Difficulty falling asleep”) and negative affect (MADRS item # 9: “Pessimistic Thoughts”). See Supplemental Table 4 for a complete list of symptom clusters and their constitutive items. Ultimately, this resolution was chosen for analysis because it both (i) allowed for interrogation of symptom dimensions finer than those aligned with traditional diagnostic categories (anxiety and depressive disorders) and (ii) resulted in communities that were broadly aligned with constructs of the RDoC, unlike finer resolutions which split groups of symptoms into clusters too fine to be meaningful in transdiagnostic analyses.

**Figure 2.**
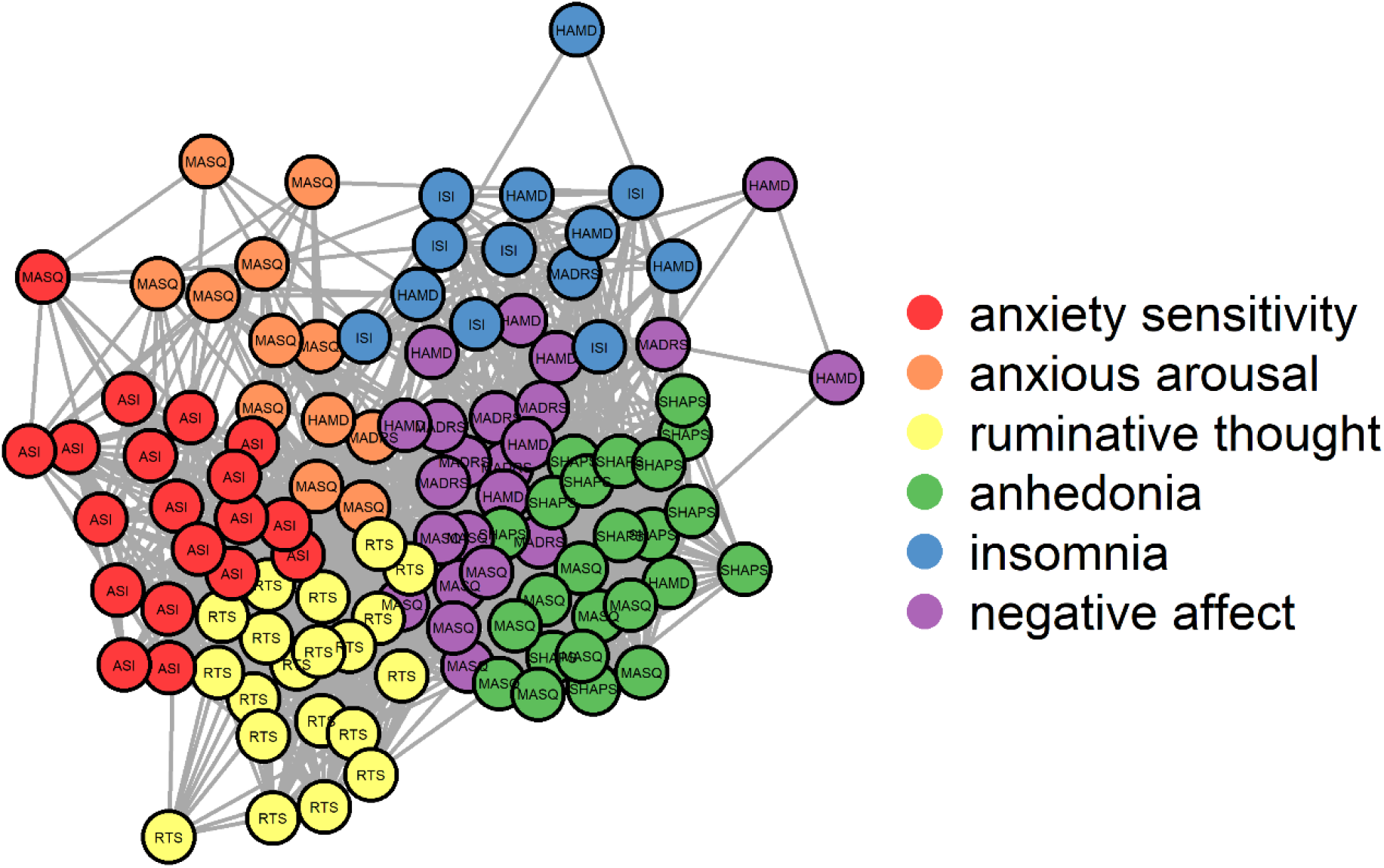
Clustering revealed six communities of symptoms. Each circle represents an item on one of seven patient assessments of psychopathology. Indicated in each circle is the assessment to which the item belongs. Connections and proximity of circles indicate the strength of correlation between items. MADRS = Montgomery-Asberg Depression Rating Scale, HAMD = Hamilton Depression Scale, MASQ = Anxiety Depression Distress Inventory-27, ASI = Anxiety Sensitivity Index, SHAPS = Snaith-Hamilton Pleasure Scale, ISI = Insomnia Severity Index, RTS = Ruminative Thought Style Questionnaire.

### Training and testing partitions

Training (n = 192) and testing (n = 50) partitions were well-matched for age, sex and *z*-scored symptom scores (Supplemental Figure 2). Two sample *t-*tests and χ2 tests did not indicate significant differences between data partitions for all categories.

### SCA results

The complete list of significant clusters identified using SCA is detailed in Table 2. Only one significant cluster was identified using a whole-brain analysis, and association with this cluster did not replicate in the testing dataset (anxious arousal, L Amygdala → whole brain, cluster region = left primary motor cortex, rtesting = 0.007, p = 0.962). Additionally, the originally proposed masks for the negative affect cluster (precuneus, mPFC and PCC) did not generate any significant clusters, so the dlPFC and insular cortices were examined post-hoc as additional masks.

Using SCA, all connectivities extracted from identified clusters were significantly correlated with symptom dimensions in the training data. However, replication within the testing data was mixed. Only clusters for anxiety sensitivity (L Amygdala → L dlPFC, rtesting = 0.337, p = 0.017) and anhedonia (R iVS → L OFC, rtesting = 0.276, p = 0.049) generated statistically significant correlations in the testing set. Other symptoms, like negative affect (sgACC → L Insula, rtesting = 0.247, p = 0.088), generated moderate, but not significant correlations, while still others, like anxious arousal (R Amygdala → R dlPFC, rtesting = 0.016, p = 0.909), failed to generate any meaningful correlations in the testing set.

### Multivariate models

#### hyperparameter tuning in training dataset

Hyperparameter tuning was accomplished using nested cross validation within the training dataset. SVR favored smaller numbers of connections (∼500 connections), with cross-validated performance (measured by Pearson’s correlation between observed and predicted symptom scores) peaking around *r* = 0.65 for most symptom dimensions (Supplemental Figure 3). In contrast, BBS favored a high number of connections (∼38000 connections), with more moderate cross-validated performance (*r* = 0.2 for most symptom dimensions). As BBS models generally exhibited increasing cross validated performance with more features, the full connectivity matrix was used for all BBS models.

**Figure 3.**
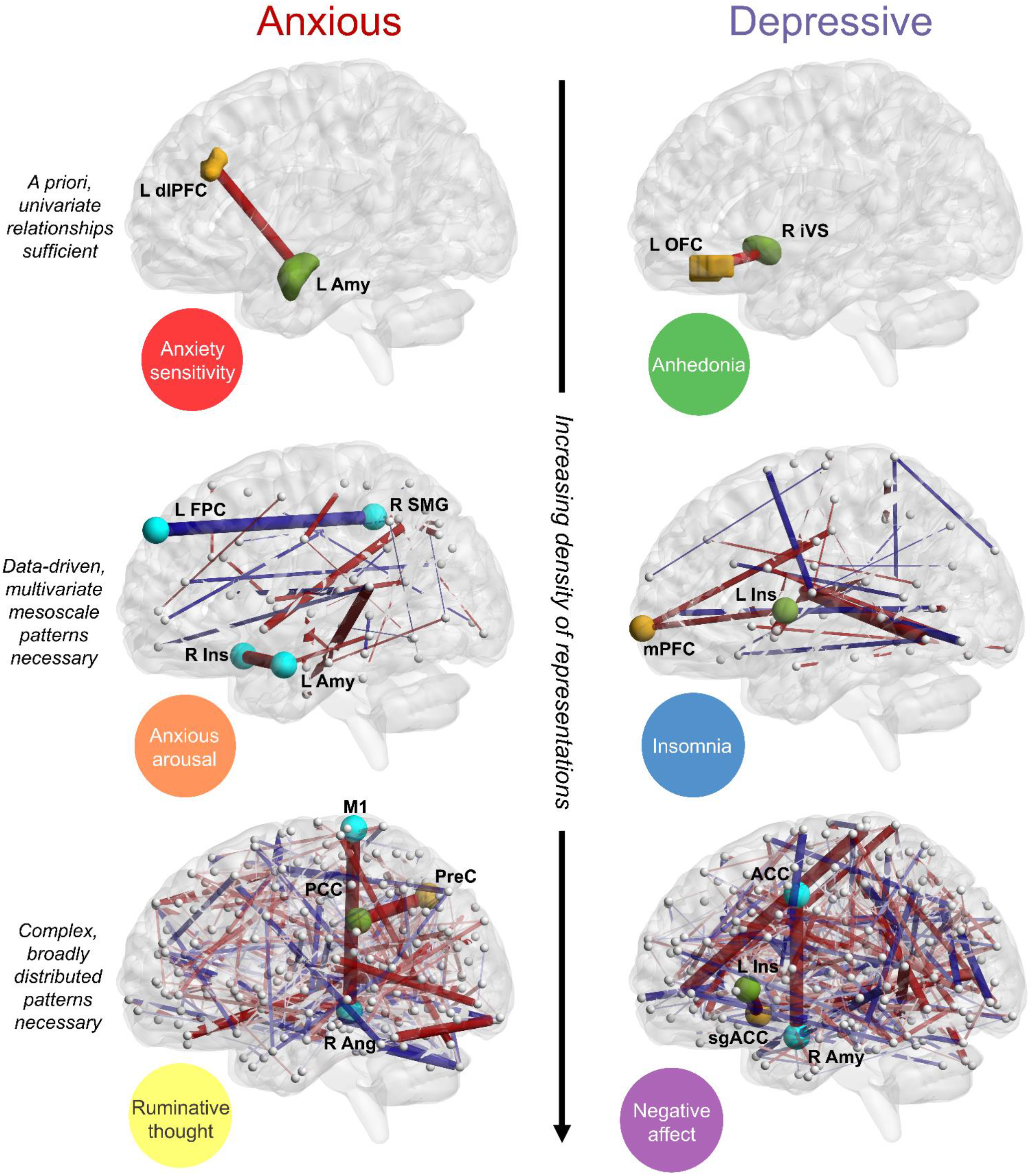
Schematic representations of each symptom and its best performing modeling approach. Symptoms in the first row were modeled using seed-based correlation analysis (SCA), in the second row using support vector regression (SVR) and in the third row using Brain Basis Set Modeling (BBS). A connection’s color represents either hyperconnectivity (red) or hypoconnectivity (blue), and a connection’s width represents the magnitude of its associated model weight. Green and yellow nodes denote the *a priori* connections tested in SCA models; green nodes represent seeds while yellow nodes represent resulting clusters. Other nodes belonging to key connections are labeled and colored cyan for easier identification: anxious arousal was associated with hypoconnectivity between L frontopolar cortex and R supramarginal gyrus and hyperconnectivity between R insula and L amygdala, ruminative thought was associated with hyperconnectivity between primary motor cortex and R angular gyrus and negative affect was associated with hyperconnectivity between R amygdala and anterior cingulate cortex. For SVR and BBS models, connections in the strongest 1% of model weights are plotted for display purposes (∼30 connections for SVR models, ∼300 for BBS models). dlPFC = dorsolateral prefrontal cortex, Amy = amygdala, FPC = frontopolar cortex, SMG = supramarginal gyrus, Ins = insula, M1 = primary motor cortex, PreC = precuneus, PCC = posterior cingulate gyrus, Ang = angular gyrus, iVS = inferior ventral striatum, OFC = orbitofrontal cortex, mPFC = medial prefrontal cortex, ACC = anterior cingulate cortex, sgACC = subgenual anterior cingulate cortex.

#### Final model performance in testing dataset

Final model performance for all three modeling approaches is detailed in Table 3. Each symptom dimension was best modeled by a different modeling approach: anxiety sensitivity (*r*testing = 0.337, p = 0.017) and anhedonia (*r*testing = 0.276, p = 0.049) were best modeled by SCA, insomnia (*r*testing = 0.344, p = 0.006) and anxious arousal (*r*testing = 0.336, p = 0.009) were best modeled by SVR and negative affect (*r*testing = 0.323, p = 0.013) and ruminative thought (*r*testing = 0.313, p = 0.014) were best modeled by BBS.

Schematic representations of each symptom and their best performing modeling approaches are displayed in Figure 3. While modeling approaches were fit independently, connections weighted heavily in simpler models were also prominent in more complex modeling approaches. For example, *a priori* connections tested in SCA generally exhibited strong weights in multivariate models: the L Insula → mPFC connection was in the top 4.04% of connections associated with insomnia, the sgACC → L Insula connection was in the top 3.50% of connections associated with negative affect and the precuneus → PCC connection was in the top 10.21% of connections associated with ruminative thought (and this connection performed comparatively worse in SCA, with *r*testing = 0.135). Only anxious arousal did not exhibit strong correspondence between its SCA model (which had poor replication in the testing set: rtesting = 0.016) and its multivariate models: SVR did not select the R Amygdala → R dlPFC connection, and this connection was only in the top 34.01% of connection weights in BBS. Finally, connection weights from SVR models correlated significantly with BBS connection weights across all symptom dimensions (*r* = 0.626, *p* < 0.001).

Network-level representations for multivariate models reveal common and distinct patterns of connectivity abnormalities associated with each symptom dimension (Fig. 4). Of the four symptoms best represented by multivariate models, all exhibited abnormal connectivity patterns in the default mode network (DMN). Anxious arousal was characterized by hyperconnectivity between bilateral regions of the DMN, as well as hyperconnectivity between (i) sensory regions like the visual and auditory cortices and (ii) networks in association cortex, like the cinguloopercular network (CIN), frontoparietal network (FPN) and DMN. Insomnia was associated with hyperconnectivity between the (i) DMN and (ii) visual and auditory cortices, the right salience network and limbic nodes. Ruminative thought was associated with hyperconnectivity between bilateral regions of the CIN, as well as hyperconnectivity between the CIN and visual cortex and hypoconnectivity between the right CIN and the DMN. Finally, negative affect was associated with decreased segregation (increased connectivity) between the left FPN and the DMN, as well as between CIN and DMN. The left FPN exhibited decreased connectivity with the salience network and visual cortex.

**Figure 4.**
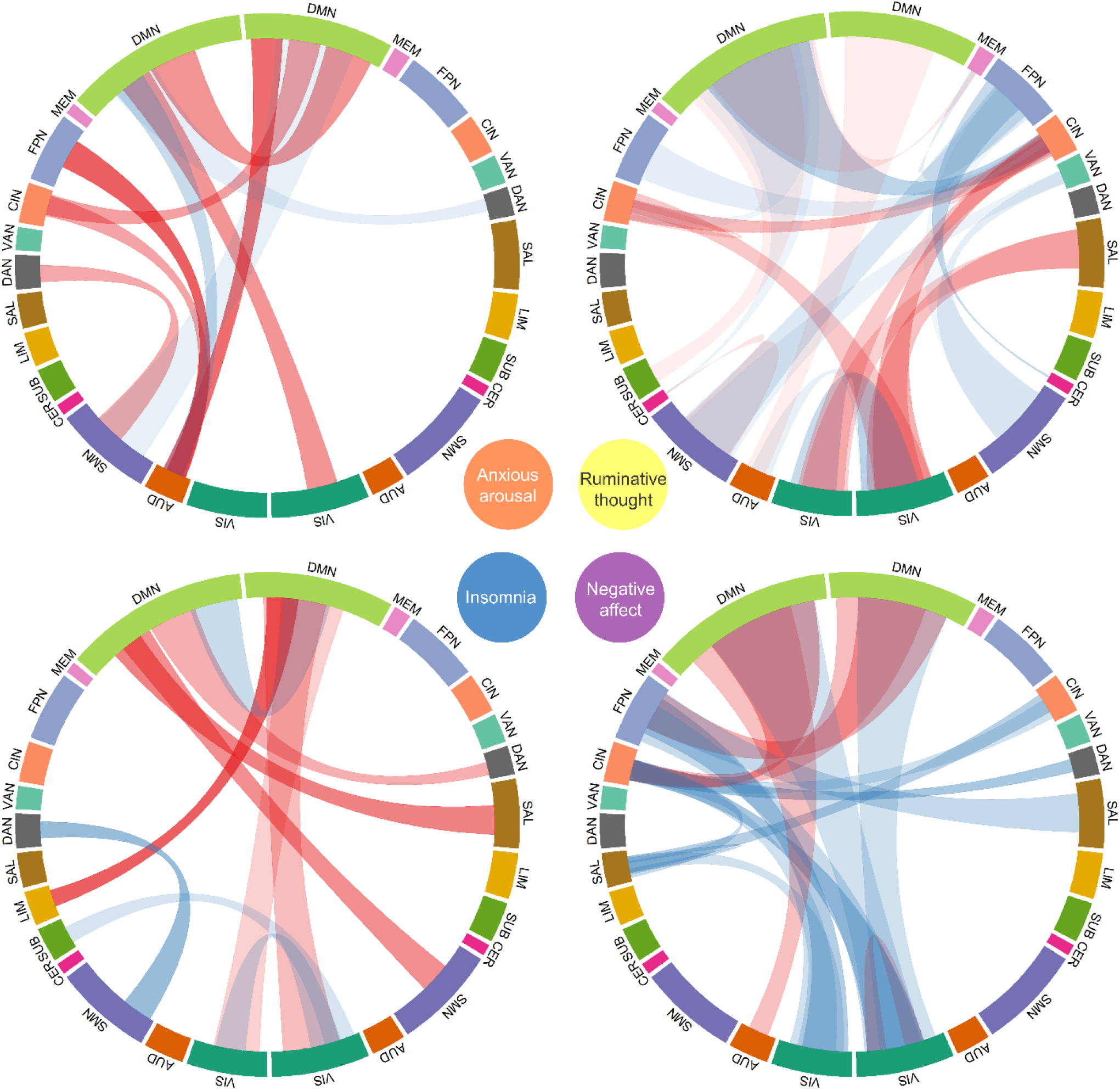
Chord diagrams displaying network-level associations for symptoms best fit by multivariate models. Association strengths were collated by node community, according to the *a priori* communities delineated in the Power264 atlas. Communities were lateralized and plotted around each chord diagram, with left-lateralized nodes displayed on the left side of the plot and bar width proportional to the number of nodes in the community. Red ribbons indicate hyperconnectivity while blue ribbons indicate hypoconnectivity in association with symptom scores. Ribbon opacity is proportional to the mean strength of association with the symptom dimension (more opaque = stronger association), and ribbon width is proportional to the number of connections participating in a plotted association (wider ribbon = more connections). Note that the left column of diagrams represents SVR models, while the right column represents BBS models. DMN = default mode network, MEM = memory retrieval, FPN = frontoparietal network, CIN = cinguloopercular network, VAN = ventral attention network, DAN = dorsal attention network, SAL = salience network, LIM = limbic, SUB = subcortical, CER = cerebellar, SMN = somatomotor network, AUD = auditory, VIS = visual. Full details for how this visualization was generated can be found in the Supplemental Methods (Under heading: *Chord diagram for multivariate models*).

## Discussion

Most studies of anxious misery pathophysiology have examined differences between a single-diagnosis cohort of patients and a matched group of healthy controls. The present work examines transdiagnostic associations between functional connectivity and different symptom dimensions, in line with the dimensional perspective espoused by NIMH’s RDoC framework [1]. Our multi-modeling approach suggests that different dimensions of psychopathology are best expressed at different scales of brain connectivity (Fig. 3). Some dimensions, like anhedonia and anxiety sensitivity, were best represented by a single, univariate relationship. Other dimensions, like insomnia and anxious arousal, required a data-driven approach, drawing from a subset of the brain connectome to generate a replicable association. Still other dimensions, like negative affect and rumination, required connectivity patterns that spanned the entire brain; for these dimensions, smaller subsets of brain connectivity measures were not sufficient to generate a replicable association.

Anhedonia and anxiety sensitivity were best modeled using single connections, suggesting that these symptom dimensions may represent more elementary neurocognitive processes gone awry. Anhedonia has been consistently linked to dysfunctions in the mesolimbic dopaminergic system, a relatively circumscribed and evolutionarily conserved system [53, 54] that plays a fundamental role in reward and goal-directed behaviors in many vertebrate species [55]. Anxiety sensitivity has been characterized as “fear of fear”, or the fear of sensations associated with the experience of anxiety [56]. Associations between anxiety sensitivity and disorders like PTSD [56, 57] and panic disorder [58], both disorders with exaggerated fear responses to threatening stimuli, support this account. Like the mesolimbic circuit in anhedonia, fear circuitry centered around the amygdala has remained well-conserved throughout evolution [59, 60].

In contrast to anhedonia and anxiety sensitivity, anxious arousal and insomnia were best modeled using SVR, which utilized a small subset of the connectome (∼5%). Like anxiety sensitivity, anxious arousal has been associated with abnormal amygdala connectivity [28, 61]. However, cross-cultural studies have revealed that different ethnoracial groups experience different degrees of anxious arousal (somatization), suggesting that this symptom dimension likely involves cortical integration beyond the amygdala [61, 62]. Indeed, anxious arousal was associated with hyperconnectivity between sensory regions and regions in association cortex (including CIN, FPN and DMN), which may underlie the somatization of anxious states. Insomnia was associated with hyperconnectivity between the DMN and multiple cortical areas, including primary sensory cortex, salience network and limbic nodes. This exaggerated connectivity with the DMN may represent increased vigilance and intrusive awareness of emotional and bodily states, resulting in poor sleep initiation and continuity [63].

Finally, two of our constructs, rumination and negative affect were best modeled by BBS, which utilized over 38,000 connections. Evolutionary theories of ruminative thought support the notion that this symptom dimension likely developed later in evolution, as the emergence of complex social environments and the ability to engage in sustained processing are thought to be supported by the expansive primate neocortex [64, 65]. Similar theories about negative affectivity suggest that stress drives the brain to reorganize to minimize surprise (prediction error) in the environment, a complex process that likely involves cortex-spanning, multi-network interactions [66]. Both ruminative thought and negative affect exhibited abnormalities with the DMN, which has been consistently implicated in anxious misery disorders, particularly in depression [41, 67]. Hyperactivity of the DMN during emotional perception and judgement [68] and passive viewing and reappraisal of negative pictures [67], as well hyperconnectivity between the DMN and the FPN [42] suggest that DMN abnormalities may underlie an inability to detach from internal emotional states (e.g., rumination and negative affect; [67]).

The present work suggests that anhedonia and anxiety sensitivity (and its related construct, fear) may be well modeled by rodent models due to high concordance with human neurobiology, while more complex symptom dimensions like negative affect and rumination may be challenging for cross-species translation. This is not to suggest, however, that rodent circuit-level analyses of negative affect and rumination are without merit; for example, the subgenual anterior cingulate cortex (sgACC) → left insula connection had a relatively robust association with negative affect, corroborating clinical neuromodulatory techniques that target the sgACC [44, 69] and rodent studies of homologous brain regions [70, 71]. In fact, across our three modeling approaches, different models of the same symptom dimension exhibited similar weights, suggesting that convergent signals informed models at different scales. However, our results do suggest that symptom dimensions differ in the degree to which they impact different scales of brain organization, and future studies in both humans and rodents should acknowledge these discrepancies.

Critical to our confidence in the validity of these conclusions is that we directly tested our models with a fully held-out validation dataset. Indeed, SVR models exhibited excellent performance in training set cross validation (typically achieving correlations of 0.6) but failed to generate significant associations in testing data for 4 out of 6 symptom dimensions. Furthermore, this gap between cross validated and testing performances persisted despite a careful segregation of feature selection across cross validation folds (Figure 2; [18]), highlighting the challenges of interpreting cross validation results without held out validation [72]. SVR models favored smaller subsets of features, typically on the order of 500 features (around ∼1% of the full connectivity matrix), which is similar to the number of connections used in many other studies examining multivariate associations of psychopathology [25, 73-75]. Notably, many of these studies did not provide a direct validation of their models on held-out data which, as the results from this study suggest, is a crucial step in assessing the generalizability of multivariate models.

While we strove to implement a rigorous analytic design and findings are reinforced by and expand upon prior literature, these results should also be considered in the context of the following limitations. In order to minimize the number of hypotheses tested for SCA, we limited seeds and masks for SCA to a circumscribed set based on extant literature. It is possible that univariate connections outside of those tested could replicate in a testing set. However, we took care to focus on those connections that were best supported by the literature. Further, exploratory whole-brain analyses only generated one cluster (anxious arousal; L Amygdala → L primary motor cortex), which failed to replicate in the testing dataset. Additionally, the testing dataset was of moderate size (n = 50); future analyses pooling data from multiple sites will prove invaluable for continuing to test the robustness and generalizability of symptom-brain associations [17].

Despite these limitations, the present work represents one of the only studies to examine transdiagnostic, multidimensional psychopathology using multiple scales of brain connectivity. Our rigorous performance assessment framework revealed that different dimensions of anxious misery psychopathology require different scales of functional connectivity, which, in addition to advancing a fundamental understanding of disease etiologies across DSM diagnosis categories, has implications for the translation of basic research paradigms to human psychopathology.

## Supplemental Methods and Materials

### Participant assessment

While participants completed a total of 32 assessments that addressed a broad range of cognitive performance, general health and social cognitive domains (full details in [1]), we focused on seven well-established scales of psychopathology that measure various aspects of anxious misery symptomology. These seven scales are: (i) Montgomery-Asberg Depression Rating Scale (MADRS[2]) and (ii) Hamilton Depression Rating Scale (HAM-D / HDRS; [3]), which both measure depressive symptoms, (iii) Anxiety Depression Distress Inventory-27 (ADDI-27 / MASQ-Short[4]), which measures both depressive and anxious symptoms, (iv) Anxiety Sensitivity Index-3 (ASI-3; [5]), which measures anxious symptoms, (v) Snaith-Hamilton Pleasure Scale (SHAPS[6]), which measures hedonic capacity, (vi) Insomnia Severity Index (ISI[7]), which measures issues with sleep and (vii) Ruminative Thought Style Questionnaire (RTSQ; [8]which measures ruminative thought severity.

### Hierarchical clustering of symptoms

To perform hierarchical clustering, we used a recently developed ensemble consensus clustering algorithm [9]), implemented in the HierarchicalConsensus package (v1.1.1) in MATLAB. We used minimum co-clustering as the measure of cohesion (option ‘min’ for ‘SimilarityFunction’) in order to prevent decreases in mean co-clustering after branching events. The result is a consensus structure representing correlational structure present in participants’ symptoms. In order to determine a reasonable resolution of symptom community structure, we manually inspected each level of the hierarchy, seeking a resolution that generated communities of the same general scale of those delineated by RDoC constructs (fear, anxiety, frustrative nonreward, sleep-wakefulness, etc.).

### Modeling approach 1

#### Seed-based correlation analysis

Seeds and masks were derived from the Harvard-Oxford Cortical and Subcortical Atlases, with supplemental regions provided by other sources when appropriate regions were not present in the Harvard-Oxford atlases (only the inferior ventral striatum and orbitofrontal cortex were from supplemental sources; see Supplemental Table 2 for a complete description of seeds and masks). In order to limit the number of hypotheses tested for SCA, each symptom dimension was associated with one set of seeds (e.g., left and right amygdalae) and one set of masks (e.g., default mode network nodes: precuneus, medial prefrontal cortex, posterior cingulate gyrus).

To conduct SCA, mean timeseries were extracted from each seed and correlated with the timeseries of every voxel in the brain, using a Pearson correlation and subsequent Fisher’s *z* transformation. Because participants completed four scans, the resulting four *z*-statistic maps were averaged within subjects to generate one map for each participant and symptom-seed. Then, linear associations with the relevant symptom dimension were tested, with significance determined by permutation testing (using FSL’s randomise, 5000 permutations; [10]

### Modeling approach 2

#### Support vector regression (SVR)

For SVR, we utilized sklearn’s (v0.23.2; Python v3.7.6) implementation, using a linear kernel [11] Tuned hyperparameters specific to this modeling approach included a regularization parameter, *C*, and the kernel coefficient, *gamma*. SVRs were fit on a subset of individual connection strengths, which were selected using our cross-validated feature selection process. Our cross-validated feature selection procedure indicated that SVR models performed best when fit using smaller numbers of connections (∼500 connections; Supplemental Figure 3).

### Modeling approach 3

#### Brain Basis Set Modeling (BBS)

BBS effectively summarizes large-scale connectivity patterns, but it is unclear whether the entire functional connectome is necessary for effective modeling. Therefore, the optimal number of connectivity features to be included in the model was determined using our cross-validated feature selection procedure; this procedure indicated that BBS models performed best when the full connectivity matrix was used (Supplemental Figure 3). Second, principal components analysis using the prcomp function in R (v3.6.3) was performed on the subjects × connections matrix (dimensions = [192 × 38503]) from the training dataset, including healthy comparators. The resulting 192 components represent orthogonal directions along which *interindividual* variance in connection strength is maximized. All 50 healthy participants were included in this modeling approach in order to capture directions of connectivity variance that extend into healthy ranges; this was the only modeling approach that utilized our sample of healthy comparators. After PCA, individual component loadings were computed by projecting each subject’s connectivity matrix onto each component. Finally, linear regression models were fit to associate each symptom dimension with these connectivity component scores.

### Feature selection for multivariate methods

With the exception of seed-based correlation analysis, which is a univariate method, all models underwent a feature selection procedure to improve performance and computational tractability. An important hyperparameter to determine was the *number of connections* to include in each model (in fact, only SVR had hyperparameters other than the number of selected connections). Feature selection was conducted for each symptom dimension that was assessed, meaning each symptom dimension was associated with a unique set of connections. Connections were selected by computing the Spearman correlations between the functional connectivity matrix and the symptom in question and then selecting the *n* connections with the highest absolute correlation. Importantly, connection selection was performed for each fold of cross-validation, within the held-in data, which is a critical step to prevent overfitting [12]

### Permutation testing for final model performance assessment

Model performance was assessed by examining the correlation between the predicted and observed symptom scores in testing set data. In order to determine the significance of these correlations, a permutation testing procedure was used wherein the rows of the training data’s imaging variables were shuffled, keeping the symptom data fixed. A model was fit on this permuted training data (rerunning nested 5-fold CV, hyperparameter tuning and feature selection each time) and then applied to the testing data, and a correlation between predicted and actual symptoms was computed. This procedure was repeated 5000 times for each symptom dimension to generate a null distribution, and a *p*-value was computed as the proportion of permuted correlations that were greater than the observed correlation.

### Chord diagram for multivariate models

In order to present the results of our multivariate models, we translated node-wise model weights to mean community-level associations (based on the *a priori* communities given by the Power atlas) and displayed these in a chord diagram (Fig. 8). For these plots, we first re-projected BBS model coefficients back into connectivity space by multiplying the vector of model coefficients with the matrix of PCA loadings. The result is a 38503 × 1 vector of loadings, one for each connection in the full connectivity matrix. Next, for both SVR and BBS models, we assigned each non-zero connection to its two communities based on the *a priori* communities given by the Power atlas, one community for each node in the connection. To further aid in interpretation and visualization, communities were lateralized such that nodes belonging to the default mode network (for example) on the left side of the brain were assigned to the left default mode community while nodes on the right side were assigned to the right default mode community, etc. Finally, visualization was accomplished by computing a “t-stat” for each community-pair, wherein the weight of each community-pair was given by the following equation:

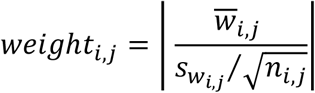

where 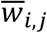 is the mean of the weights assigned to connections in community-pair 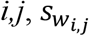 is the standard deviation of those weights and *nij*is the number of connections constituting community-pair *i,j*. The sign and absolute value of this term determined the color and opacity of each connection, respectively, in the resulting chord diagram (Fig. 8). The width of ribbons was set to be proportional to the number of constitutive connections. Community-pairs were plotted until the cumulative *weight* reached a specific threshold (150 for BBS models, 100 for SVR models), a threshold which was selected to balance interpretability and faithful representation of model complexity.

**Supplementary Table 1.**
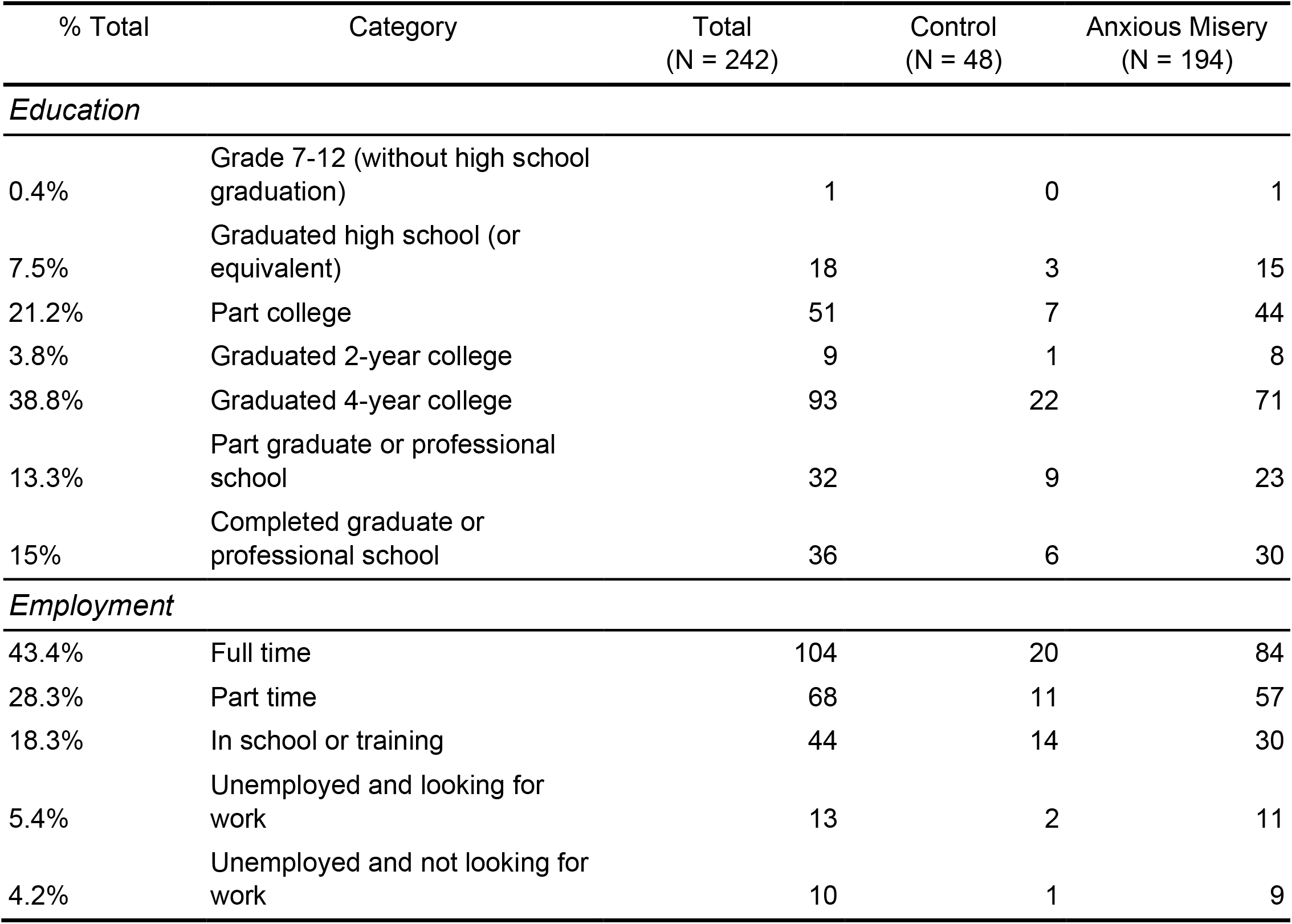
Additional Demographic Characteristics

**Supplementary Table 2.** Seeds and masks used for SCA analysis

| Region name | Atlas | Name in atlas | MNI coordinates<br>(x,y,z, radius) |
| --- | --- | --- | --- |
| L/R Amygdala | Harvard-Oxford | L/R Amygdala | -- |
| L/R dlPFC | Harvard-Oxford | L/R Middle Frontal Gyrus | -- |
| PCC | Harvard-Oxford | Cingulate Gyrus, posterior division | -- |
| Precuneus | Harvard-Oxford | Precuneous Cortex | -- |
| mPFC | Harvard-Oxford | Frontal Medial Cortex + Frontal Pole | -- |
| iVS* | -- | -- | (±14, 8, -9, 3) [13] |
| OFC | AAL | Superior frontal gyrus, orbital | -- |
| sgACC | Harvard-Oxford | Subcallosal Cortex | -- |
| Insula | Harvard-Oxford | Insular Cortex | -- |

Abbreviations: Harvard-Oxford = Harvard-Oxford Cortical and Subcortical Atlases, AAL = Automated Anatomical Labeling Single-Subject Atlas, dlPFC = dorsolateral prefrontal cortex, PCC = posterior cingulate gyrus, mPFC = medial prefrontal cortex, iVS = inferior ventral striatum, OFC = orbitofrontal cortex, sgACC = subgenual anterior cingulate gyrus. \* All regions used in SCA analysis were drawn from atlases, with the exception of the iVS seed (as neither Harvard-Oxford nor AAL atlases had sufficiently fine parcellations of the striatum). For this seed, we used spherical ROIs centered at the listed MNI coordinates.

**Supplementary Table 3.** Additional ROIs related to anxious misery pathophysiology added to the Power264 atlas

| Name | x | y | z |
| --- | --- | --- | --- |
| L nucleus accumbens | -13 | 8 | -8 |
| R nucleus accumbens | 13 | 8 | -8 |
| L subgenual anterior cingulate | -5 | 15 | -11 |
| R subgenual anterior cingulate | 5 | 15 | -11 |
| L caudate nucleus | -12 | 18 | -3 |
| R caudate nucleus | 12 | 18 | -3 |
| L amygdala | -19 | -2 | -21 |
| R amygdala | 19 | -2 | -21 |
| L hippocampus | -26 | -12 | -22 |
| R hippocampus | 26 | -12 | -22 |
| L locus coeruleus | -5 | -37 | -27 |
| R locus coeruleus | 5 | -37 | -27 |
| ventral tegmental area | 6 | -15 | -15 |
| raphe nucleus | 6 | 24 | -16 |

All coordinates are given in MNI coordinate space. ROIs were generated using 5mm radius spheres centered on the coordinates.

**Supplementary Figure 1.**
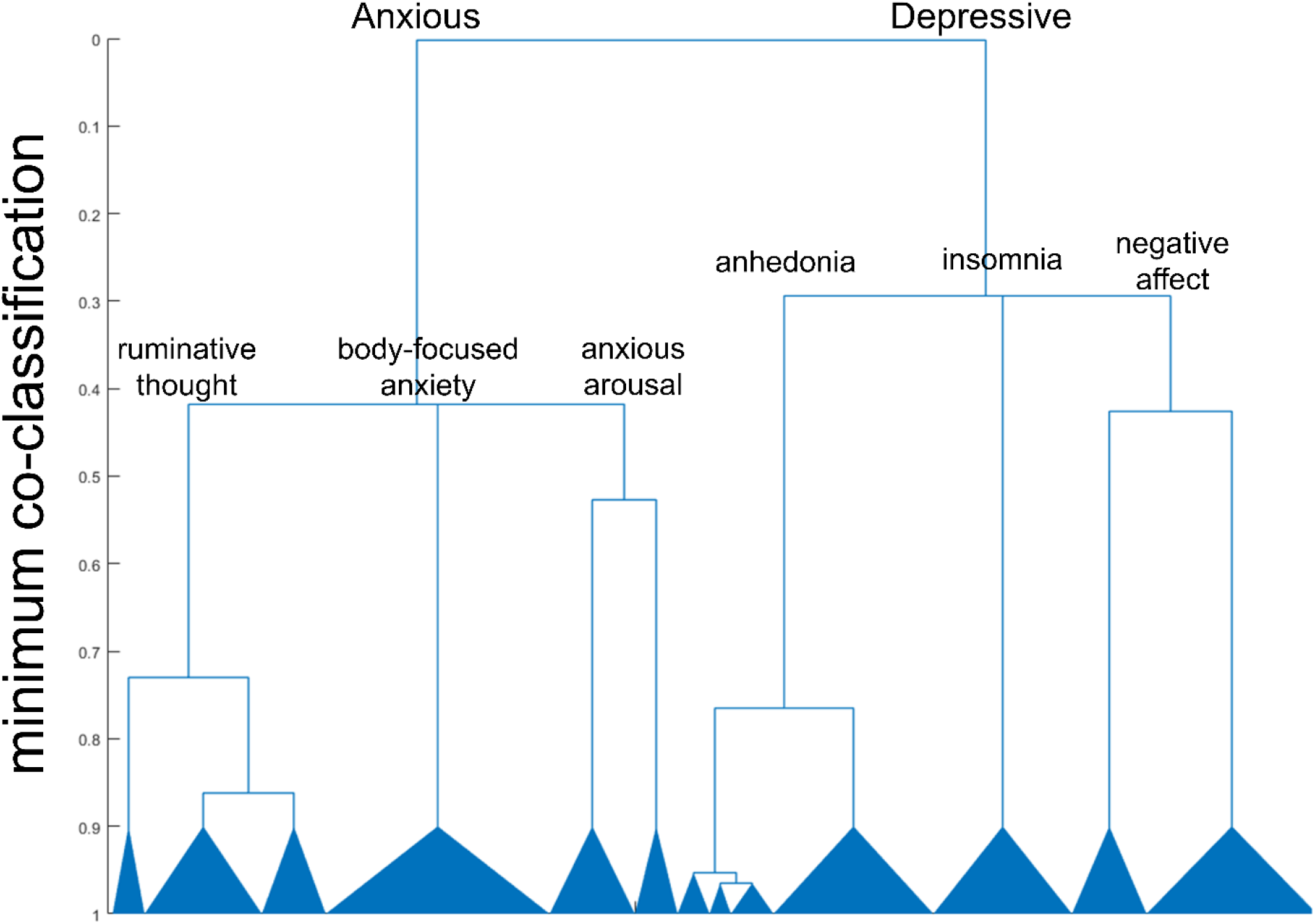
Consensus hierarchical clustering result of participant symptoms. Labels were generated through manual inspection of symptom clusters. Minimum co-classification was selected as the measure of cohesion, and 1000 partitions where each partition is sampled using a different value of γ) were used. Figure was generated using the drawHierarchy function of the HierarchicalConsensus package in MATLAB, v1.1.1 [9]

**Supplementary Table 4.**
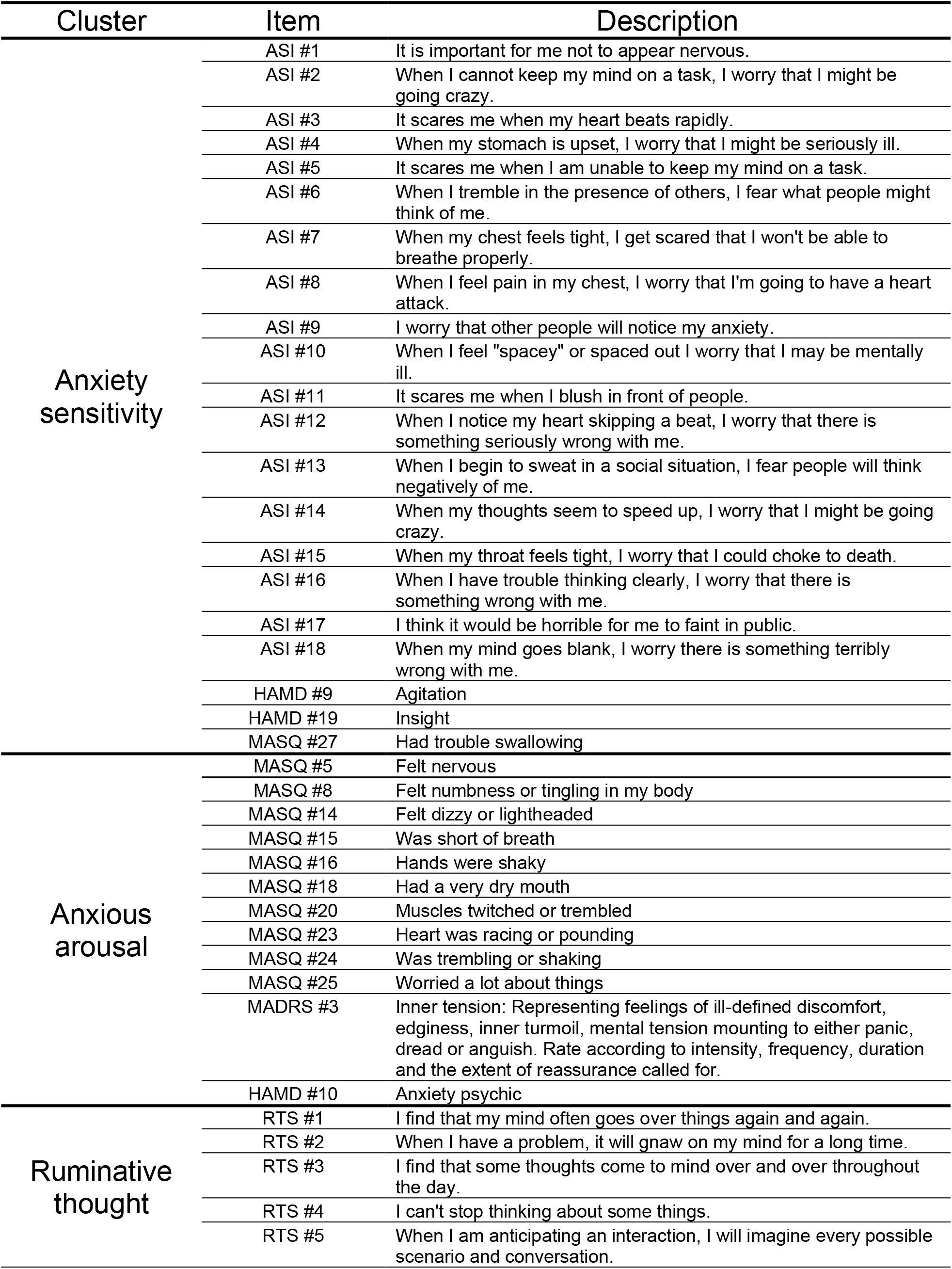
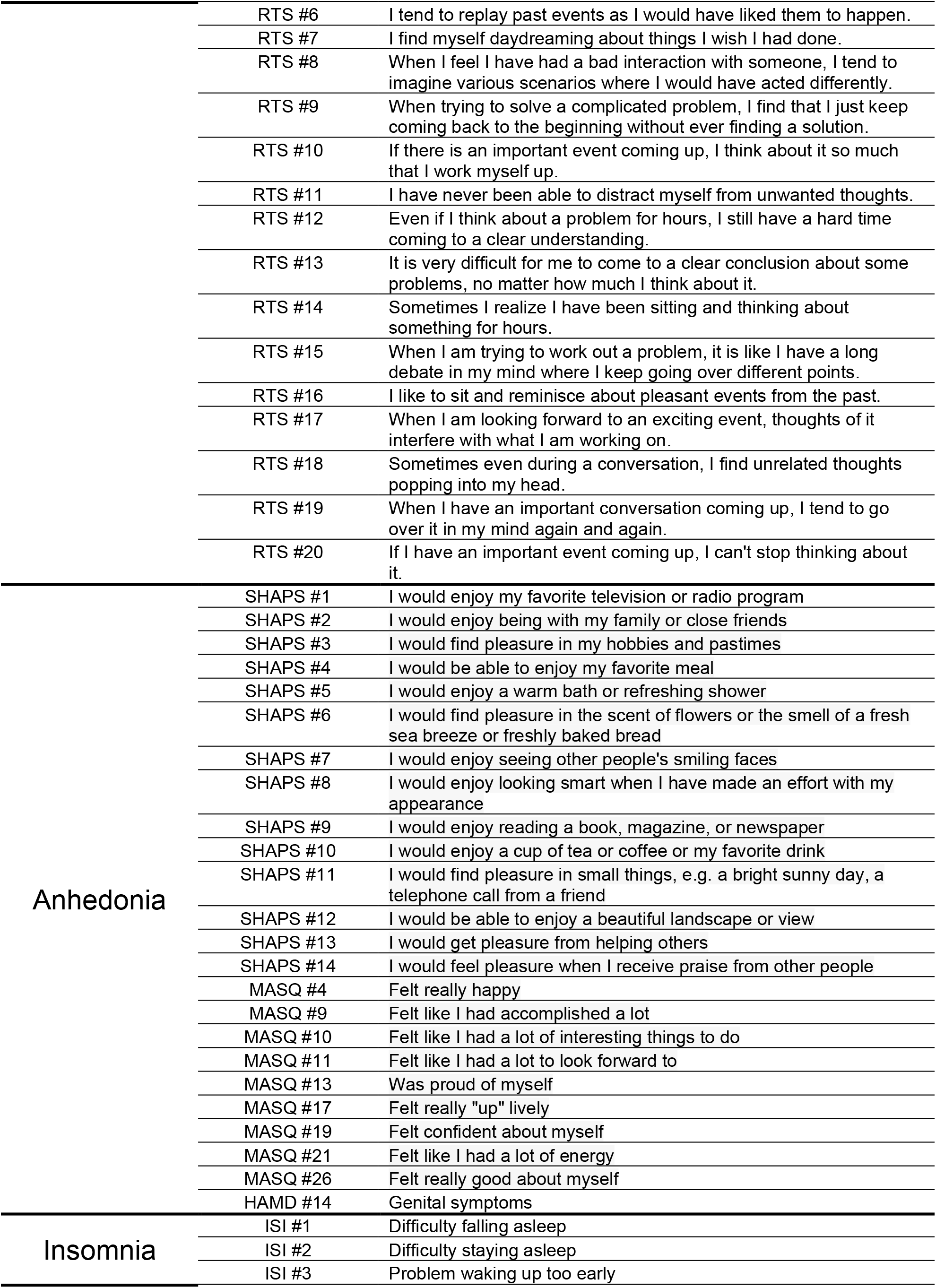
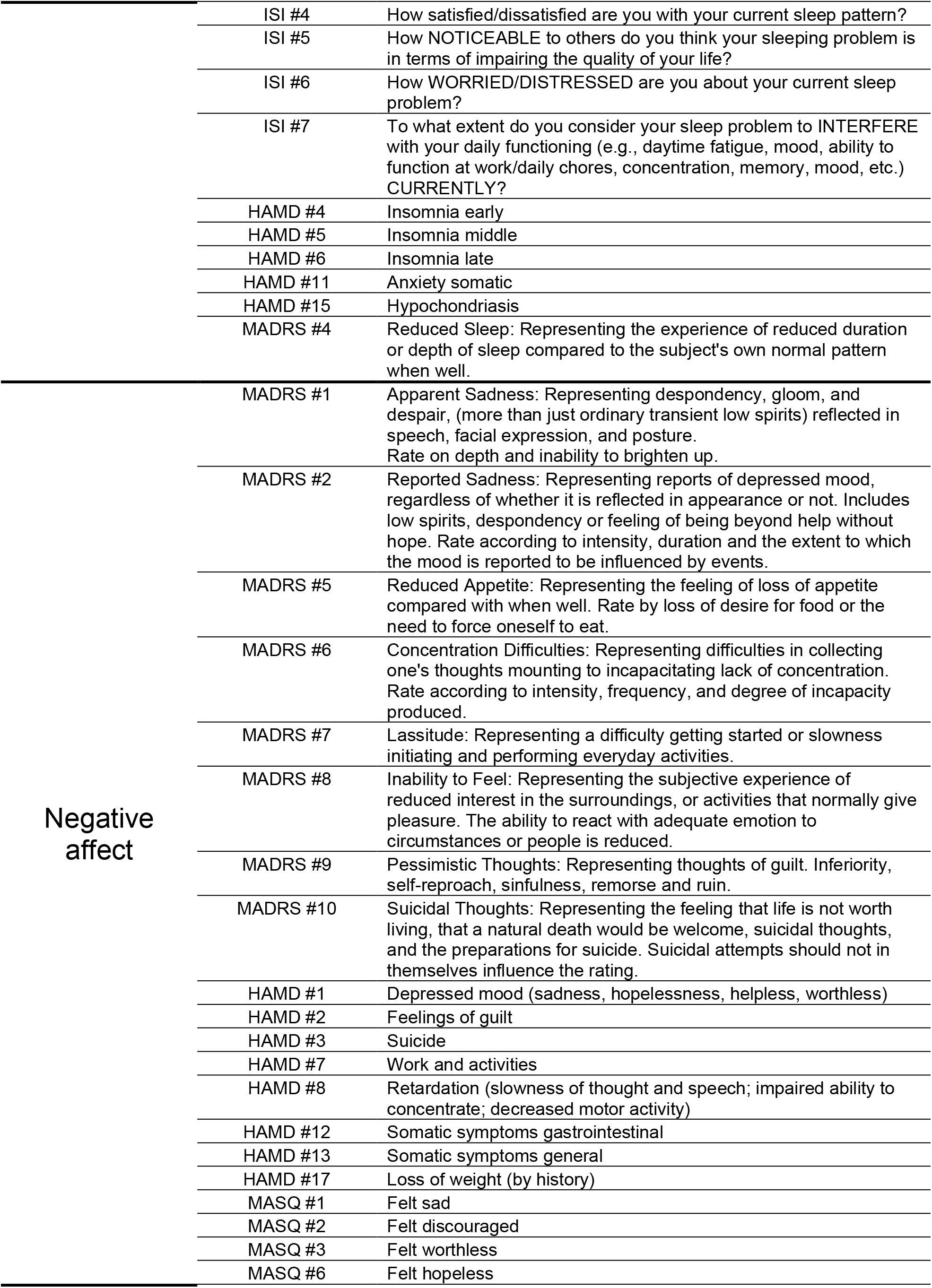
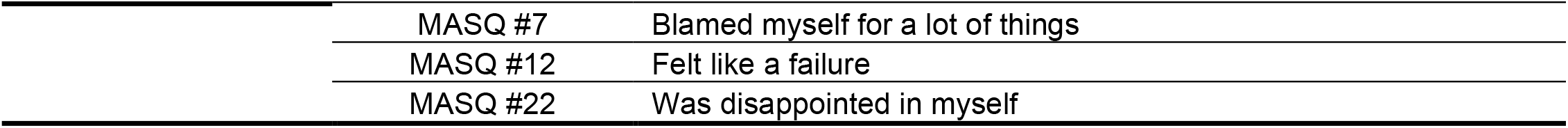
Complete list of symptom clusters and constitutive items

**Supplementary Figure 2.**
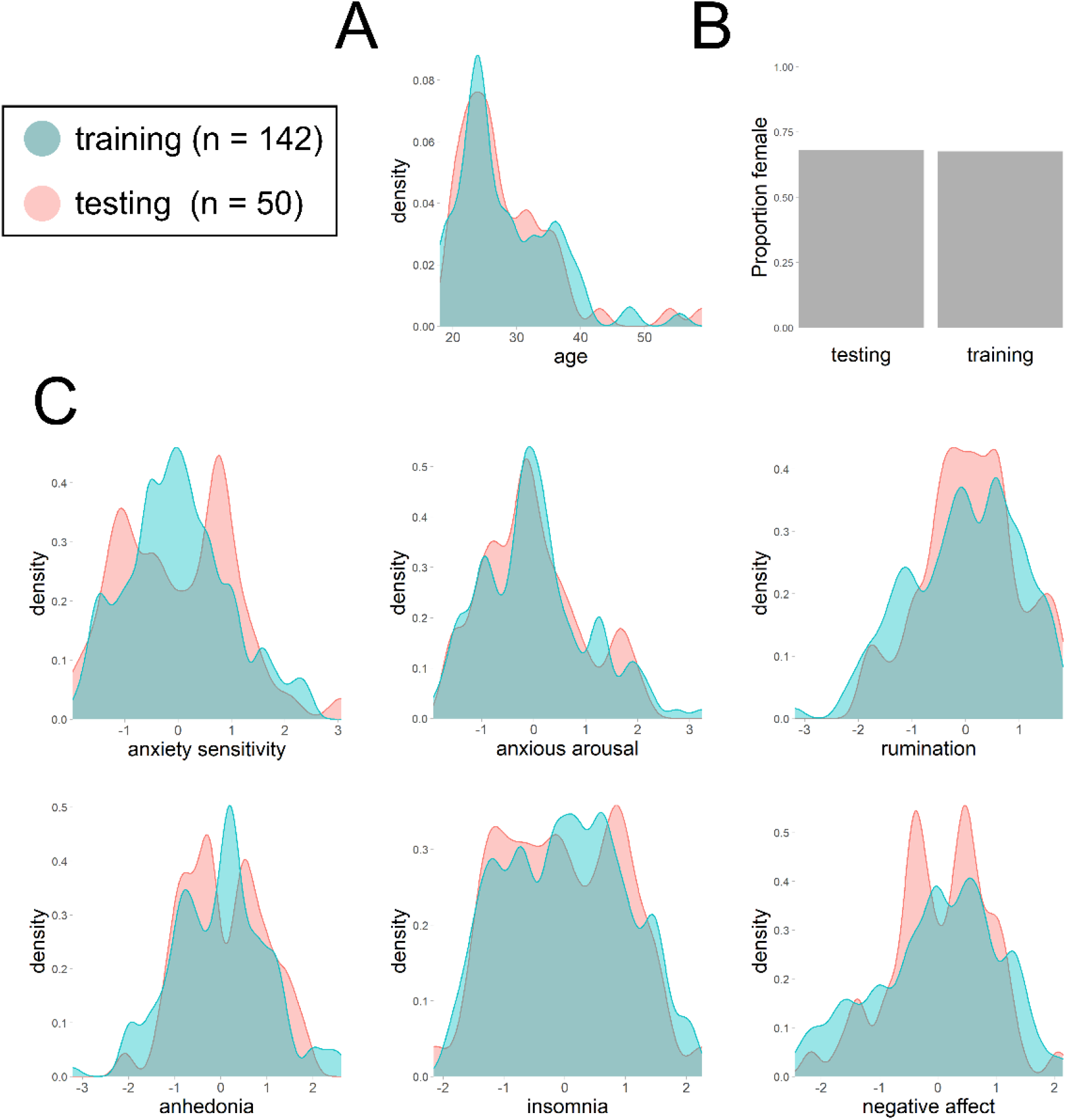
Training and testing partitions were matched for age, sex and symptom severity. **A**. Density plot of age. **B**. Barplots of proportion female. **C**. Density plots for *z-*scored symptom dimensions, one for each of our six dimensions.

**Supplementary Figure 3.**
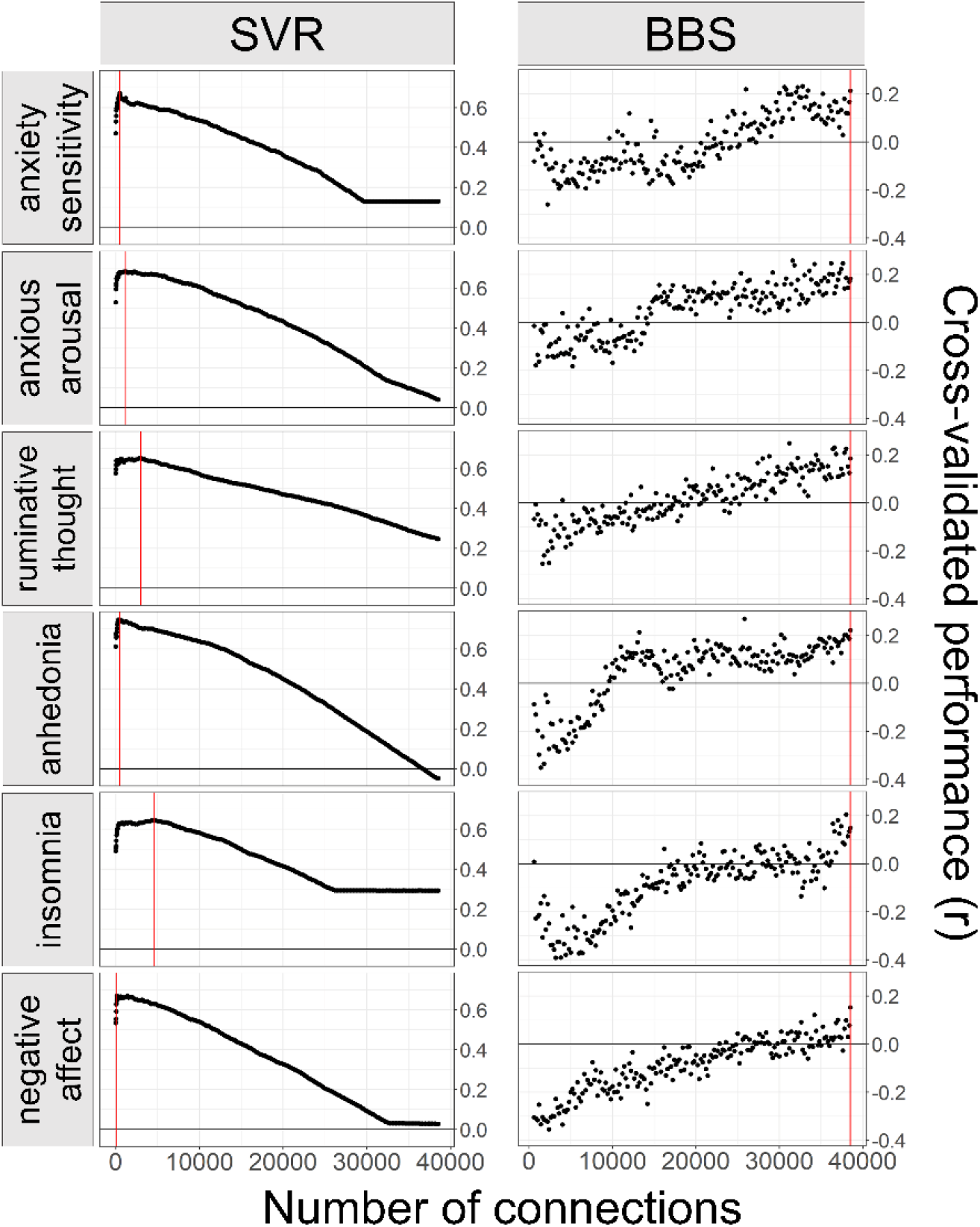
Results from cross-validation identified the optimal number of connections to select for each method and symptom dimension. Y-axis indicates the mean Pearson’s correlation between predicted and observed symptom score across a 5-fold cross validation scheme. Red line indicates the number of connections used in the final model fit on the full training dataset. Cross-validation performance for BBS exhibited increasing performance with more connections, so the full connectivity matrix was used for all BBS models. SVR = support vector regression, BBS = Brain Basis Set modeling.

